# Screening Envelope Genes Across Primate Genomes Reveals Evolution and Diversity Patterns of Endogenous Retroviruses

**DOI:** 10.1101/2024.10.28.620668

**Authors:** Saili Chabukswar, Nicole Grandi, Elena Soddu, Liang-Tzung Lin, Enzo Tramontano

## Abstract

Endogenous Retroviruses (ERVs) are integrated into the host DNA as result of ancient germ line infections, majorly by now extinct exogenous retroviruses. Accordingly, vertebrates’ genomes contain thousands of ERV copies, providing “fossil” records for the ancestral retroviral diversity and its evolution within the host. Like exogenous retroviruses, ERV proviral sequence consists of *gag, pro-pol*, and *env* genes flanked by long terminal repeats (LTRs). In particular, the characterization of *env* gene changes over time allows both to understand ERVs evolutionary trajectory and to infer their potential role in host physiology, including their pathological involvement. Despite their potential impact on the host, a detailed characterization has been performed for few domesticated *env* genes, while a comprehensive survey of their abundance and diversity among primates’ genomes is still lacking. The study primarily focused on ERVs’ *env* that are known to be present in hominoid species, given their well-characterized nature and availability in public databases. Hence using these *env* sequences, we reconstructed 32 Env sequences representing the prototypes of these ancestral proteins in Class I, Class II, and Class III HERVs. These reconstructed Envs were then used for similarity search, phylogenetic analysis, and examination of recombination events occurred within primates’ genomes that were applied to 55 primate species across the *Catarrhini* and *Platyrrhini* parvorders. Through such comprehensive pipeline, we reconstituted a phylogenetic distribution of ERV based specifically on the *env* genes, showing that the ERVs have been prevalent and widely distributed across the primate lineage. We observed for the first time the presence of the HML (human mouse mammary tumor virus-like) group in the *Platyrrhini* parvorder, indicating initiation of spread of HML supergroup before the split between New World Monkeys (NWM) and Old World Monkeys (OWM) i.e. even before 40 mya. Importantly, we confirmed interclass and intra-class *env* recombination events confirming the phenomenon of “*env* snatching” among primates’ ERVs. As a result, we demonstrate that tracing the diversity patterns of ERVs’ *env* provides relevant insights into the retroviral evolutionary history of ERVs in *Catarrhini* and *Platyrrhini* parvorders. Overall, our findings reveal that *env* recombination contributes to the diversification of ERVs, thereby broadening our comprehension of retroviral and primate evolution.

## Introduction

Retroviruses, discovered over a century ago, are a diverse group of animal viruses characterized by a common replication mechanism (Mager & Stoye, 2015). Their lifecycle involves reverse transcription of a single-stranded RNA (ssRNA) into a double stranded DNA (dsDNA), which integrates into the host genome forming a provirus (Mager & Stoye, 2015). While the retroviruses circulating nowadays typically infect somatic cells, an abundance of ancient retroviruses have been able to integrate their genome into germline cells, becoming endogenous retroviruses (ERVs) and allowing their vertical transmission from one host to another through generations (Sverdlov, 2000). Over the span of millions of years, thousands of ERV copies have been accumulated throughout the vertebrate lineage and can range from complete proviral structures to highly fragmented remnants of the provirus (Hayward, 2017; Mager & Stoye, 2015; Weiss, 2006).

Likewise exogenous RVs, ERVs also consists of four major viral protein-coding genes i.e. *gag, pro-pol* and *env* flanked by identical long-terminal repeats (LTRs) at 5’ and 3’ ends of the provirus (Hayward, 2017; Johnson, 2019). Among the four viral genes *gag* and *pro*-*pol* are well conserved across retroviral species while *env* is the most diverse, being responsible for the specific tropism of the infection (Henzy & Johnson, 2013). Given that the reverse transcriptase (RT) domain of the *pol* gene is a highly conserved region, it is widely used to infer phylogenetic relationships and to categorize retroviruses into distinct taxonomic lineages within the Retroviridae family (Henzy & Johnson, 2013). Based on RT sequences, ERVs are classified into three main classes i.e. Class I (Gammaretroviruses and Epsilonretroviruses), Class II (related to Deltaretroviruses, Lentiviruses, Betaretroviruses, and Alpharetroviruses) and Class III (Spumaretroviruses) (Blomberg et al., 2009; Gifford et al., 2018). Various phylogenetic relationships of ERVs are also based on the comparisons between RT and the transmembrane (TM) region of the *env* gene, however discrepancies may arise due to the diversity of the *env* gene among different ERVs of the same class (Blomberg et al., 2009). While the analysis of RT sequences identifies the evolutionary lineage that gives rise to a retroviral species, studying the *env* gene provides insights into the bases of such retroviral evolution, signaling the emergence of new viruses leading to cross species transmission.

The ERVs’ Envelope (Env) glycoprotein encoded by *env* gene consists of two main components, i.e. surface (SU) and transmembrane (TM) units, which includes some domains used for its characterization and phylogenetic analysis: the immunosuppressive domain (ISD), the fusion peptide (FP), the heptad repeats (HR1 and HR2), and the CX(6)C motif(Chabukswar et al., 2023). In addition, the most recent ERV group, the HML2, is divided into type I and type II due to the presence of alternative splicing variants of *env* thus producing Np9 (type I) and Rec (type II), differing in the presence and absence of a 292 nucleotide (nt) deletion in the *env* gene, respectively (Chabukswar et al., 2022; Heyne et al., 2015; Subramanian et al., 2011). The divergence in the *env* gene can be accounted by two main factors i.e. mutations and recombination. Mutations lead to modification in *env* nt sequence, whereas recombination events involve the transfer of *env* portions across viral genomes. Recombination occurs between viral genomes present in the same host cell as they exchange one or more genetic portion(s), potentially leading to the emergence of new viral strains and viruses, expansion of viral lineages, increase in their virulence and pathogenicity, and resistance to antiviral drugs (Pérez-Losada et al., 2015). A high degree of mosaicism in the *env* gene has been previously reported in the form of recombinations or secondary integrations, and tends to complicate the sequence-based HERV classification (Vargiu et al., 2016). Similarly, we previously observed the presence of recombinant forms of HML2 elements in Old World Monkeys (OWMs), specifically in *Macaca mulatta* and *Macaca fascicularis* (Chabukswar et al., 2022).

The analysis of the patterns of *env* diversity is an essential step to better understand the evolution of ERVs and reconstruct the structure of the ancestral ERV viral genes, which in turn can help in various functional analysis of the correspondent ancestral proteins, as the changes in the genes overtime has affected their production and functions. Hence, in the present study we aimed to reconstruct the *env* gene from the different HERV groups by generating a representative collection of 32 full-length consensus Envs for Class I, Class II and Class III. These proteins were used for the screening of representative species of *Catarrhini* (*Hominidae, Cercopithecinae* and *Colobinae*) and *Platyrrhini* parvorders, identifying *env* genes and investigating their diversity by implementing a similarity search strategy, followed by the study of their phylogeny and recombination patterns. With this approach, we provided a comprehensive description of the ERV *env* diversity in 55 primates species, showing the presence of several class II ERVs in the *Platyrrhini* parvorder that have not been previously reported, and detailing the occurrence of events of interclass and intraclass *env* recombination that significantly expand our understanding of retroviral evolution.

## Methods

### Generation of Representative HERV Envelope protein sequences

The *env* consensus sequences that have been previously reported in Table 4 and Table 6 of Vargiu et al. 2016 were collected and used as a query to extract the correspondent *env* genes for each HERV group from human genome (GRCh38/hg38) available on UCSC Genome Browser through the BLAT search function. To build these consensus sequences, we followed different approaches, we first extracted a total of 33 HERV elements that have been previously reported in Table 6 of Vargiu et al.2016, as the ones including the most intact *env* genes, belonging to the Class I groups HERV9, HERVFC, HERVH, HERVT, HERVW, HERVE, HERV3 and the Class II groups HML2 (18 out of 33 sequences), HML5, HML6, and HML8 For the other HERV groups not included in this list, we referred to the Table 4 of Vargiu et al., 2016, which provides a summary of 39 canonical HERV clades found in the human genome (GRCh37/hg19). More 190 sequences were extracted from the additional files of Vargiu et al., 2016 for which we generated a bed file and extracted them with the flanking of 200 nt at both 3’ and 5’ ends using the table browser tool in the genome browser (Supplementary file1 Sheet1 (S1)). These 190 sequences initially extracted were divided into canonical and non-canonical forms (Vargiu et al., 2016) where canonical refers to sequences that are well-preserved and match the structural and functional features of complete *env* genes, and non-canonical refers to sequences with significant structural alterations or truncations that deviate from this typical form (Vargiu et al., 2016). These sequences were further aligned group-specifically, to understand if any of the sequence codes for complete Env proteins. Further, to select some sequences to be used as a reference, we extracted 19 consensus sequences from Dfam (both nt and aa), and 17 reported HERV-Env proteins from UniProt for which the accession numbers are provided in Supplementary file 1 Sheet2 (S2). The sequences used to perform alignments for each HERV group are described in detail in Supplementary file 1 Sheet3 (S3), which were then aligned to references sequences taken from Dfam, NCBI and UniProt sequences (Supplementary file 1 (S2)).

For each HERV group alignment, we translated the nt sequences in 3 forward frames to check the presence of stop codons using the Geneious software (http://www.geneious.com). Furthermore, in order to reconstruct the Env proteins, we translated *env* nt sequence alignments into all three forward frames in Geneious to identify the translatable regions. The correct reading frame was identified according to the alignment and comparison with known Env domain and the high sequence similarity to the reference Env proteins taken from Dfam and Vargiu et al. 2016. The insertions or deletions (indels) were retained in the initial alignment but were carefully managed during the reconstruction process. Regions with ambiguous or excessive indels that compromised the alignment integrity were further masked or excluded. The correct reading frame was further validated by identifying the conserved domains characteristic of Env proteins, such as signal peptide, SU, and TM domains using NCBI CD prediction tool. The frames lacking these hallmark features were excluded from the reconstruction process. The Env protein consensus sequences were generated for each group by realigning the extracted translated portions to the reference sequence to generate a complete representative Env sequence for each HERV group. These generated Env sequences were then aligned to the exogenous retroviruses of all classes. For this, we collected 52 envelope protein sequences from UniProt database that included different strains as well as subtypes of exogenous retroviruses from all the genera (i.e. alpha, beta, gamma, delta, epsilon, intermediate-Beta like, spuma and lentiviruses). The Uniprot accession numbers for the Envs of the exogenous retroviruses have been provided in Supplementary file 1 (S2).

### ERV Env Screening in Primate Genomes

The reconstructed HERV Env sequences were used as a query to analyze 55 primate genomes, including the representative species from both the *Catarrhini* (*Hominidae, Cercopithecinae* and *Colobinae* families) and *Platyrrhini* parvorders. All the genome assemblies were sourced from NCBI Assembly database (https://www.ncbi.nlm.nih.gov/assembly/). The screening of the genomes for the *env* sequences was done by performing the tBLASTn (https://blast.ncbi.nlm.nih.gov/Blast.cgi) search for all the HERV groups separately. The tBLASTn algorithm uses amino acid sequences to compare the translated nt database in all six frames, and hence the nt hits obtained were further refined. The search parameters included a minimum sequence identity threshold of 40% and an E-value cutoff of 1E-5. Hits shorter than 40% of the complete *env* gene were excluded from further analysis. To ensure specificity, secondary integrations and repetitive elements were manually curated and removed. All the primate genome assemblies that were screened during the tBLASTn search are listed in the supplementary file 2 while the accession numbers for hits obtained for gamma-like, beta-like and spuma-like are available in the Supplementary phylogenetic figures 3, 4, and 5.

### Sequence Alignments and Phylogenetic Analysis

All the hits obtained from the tBLASTn search were filtered and were downloaded in fasta format. For each group, sequences whose length was less than 50% of the entire *env* gene were discarded. We then performed group specific nt sequence alignments using MAFFT algorithm of Geneious software. These alignments were further used to carry out phylogenetic analysis with the IQTREE software (http://www.iqtree.org/) for gamma-like, beta-like and spuma-like *env* nt sequences. The trees were inferred using maximum likelihood (ML) methods in IQ-TREE with a bootstrap of 1000 replicates. The trees generated were further visualized and annotated by the Figtree V1.4.3. software(http://tree.bio.ed.ac.uk/software/figtree/).

### Detection of Recombination Events

After the phylogenetic analysis of gamma, beta and spuma-like ERVs, the recombination events were detected using RDP, Geneconv, Bootscan, Maxchi, Chimaera, Siscan and 3SEQ methods implemented in RDP4 software (http://web.cbio.uct.ac.za/~darren/rdp.html) (Martin & Rybicki, 2000). The default settings of the RDP4 software that were used are (i) only potential recombination events detected by four or more of the above-mentioned methods; (ii) sequences were treated as linear and (iii) window size of 30 variable nt positions used for the RDP method. The breakpoint positions and the recombinant sequences indicated for the potential recombination event were further confirmed through comparative genomics. The detected recombinant sequences were further used to perform BLAT search in the primate genomes available in UCSC Genome to confirm the position and location of recombinant sequence in the host genomes. To highlight the initiation of the recombination events, a phylogenetic tree of all the genomes of *Catarrhini* parvorder available on the genome browser was generated using the TimeTree server by simply uploading the list of primate species on the server. The tree reflects host phylogeny and was based on whole-genome sequences, except for Gorilla species, since the *Gorilla gorilla gorilla* genome was used to generate the primate phylogeny. We further annotated the events on the phylogenetic tree based on the genomes in which the recombinants were detected.

## Results

### Reconstruction of the Retroviral Envelope from Human ERVs

To understand the ERVs evolutionary dynamics, reconstructing the ancestral ERV prototypes is a crucial step and requires to address two important issues i.e., the biases present in the ERV data used and the changes that had occurred after each single retrovirus was integrated into the host genome. With the purpose of providing prototype Env proteins for the different HERVs, we reconstructed the Env sequence of 32 HERV groups, including members of Class I (gamma- and epsilon-like, 20), II (beta-like, 10), and III (spuma-like, 2) (Table1, Supplementary file1(S4)). To ensure accuracy, all reconstructed sequences were manually curated to verify the presence of functional regions and sequence integrity. A schematic representation of the Env reconstruction process is shown in Figure 1.

**Figure 1:**
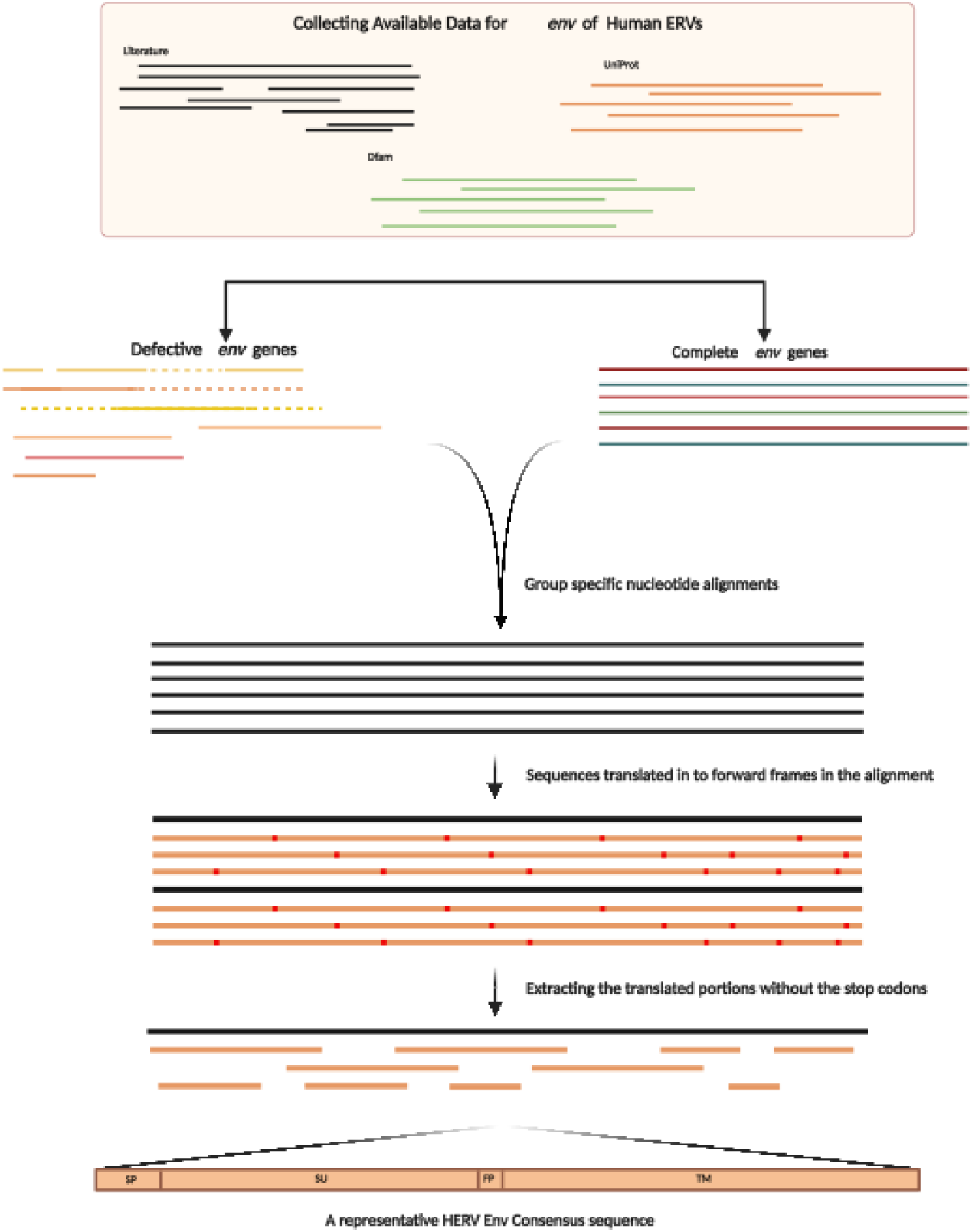
Illustration of the reconstruction process of HERV Env proteins. The first step was to extract *env* genes for both endogenous and exogenous retroviruses from Literature, Dfam, NCBI, Uniprot, and PDB (complete *env* sequence as well as defective *env* genes). Second, we performed group specific alignments for all the ERV sequences with Mafft and further divided the sequences into subtypes based on the alignments, as in case of HERV3 and HML3. Then, we translated the nt sequences into three forward frames in the alignment and extracted only the translated portions that did not have any stop codons. Lastly, the extracted aa portions were aligned to the reference sequence of each group and hence, a reconstructed env sequence was generated for HERV groups as mentioned in Table 1.

**Table 1:**
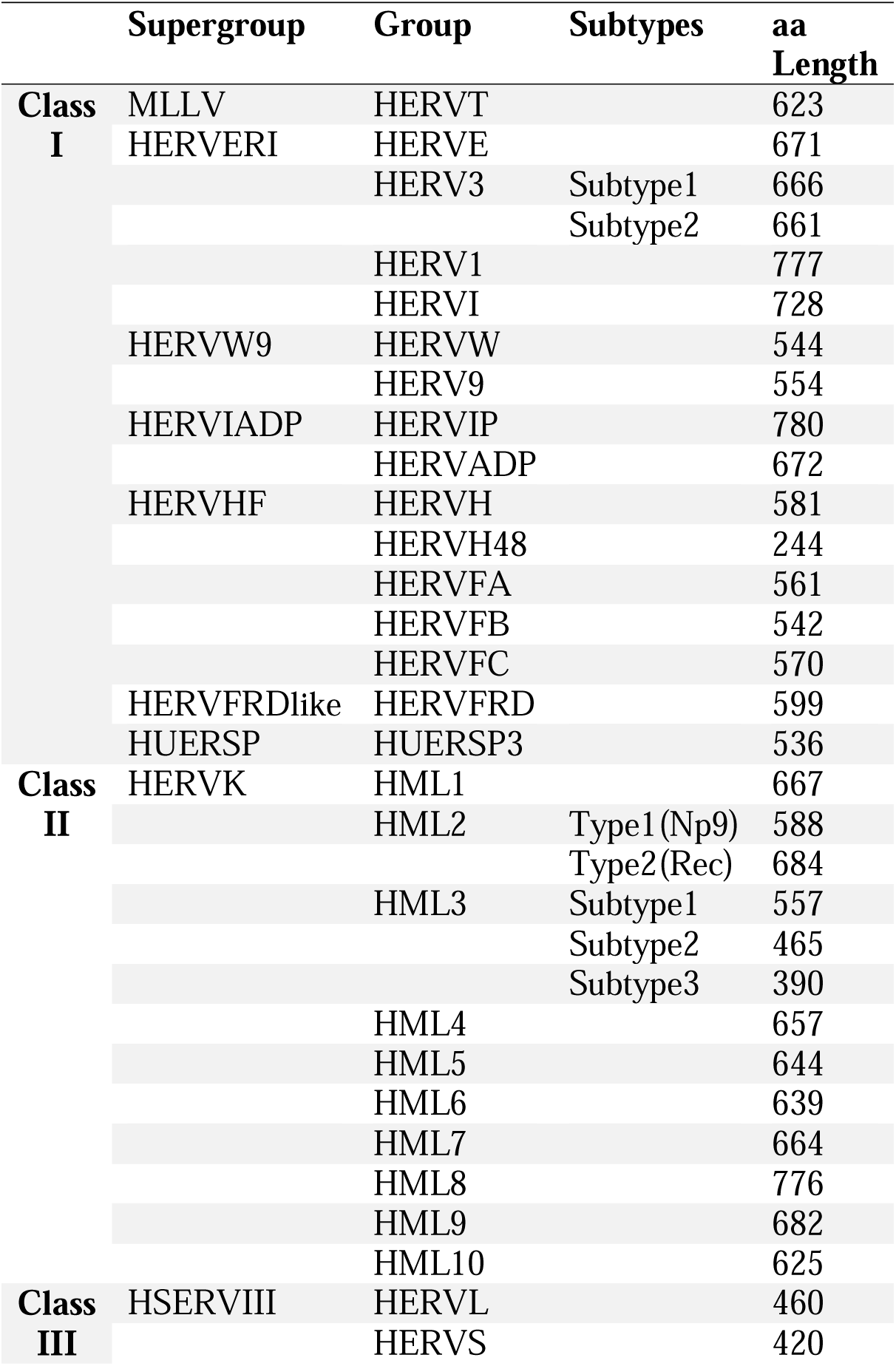
Dataset of the reconstructed HERV Envelope proteins.

Since the Env reconstruction was performed based on the data available for HERVs, to confirm the accuracy of reconstruction by its relatedness to the other RVs, we generated Env protein phylogeny of reconstructed Envs with 58 Env proteins of exogenous RVs (Figure 2). As expected, we observed a class specific clustering of the reconstructed Envs with that of exogenous RVs, except for HERVS which was clustered with Class I RVs and not with the Class III RVs, indicating that HERVS has a Class I Env that is related to PRIMA41, PABL and HERV1_Artodact, as also previously reported (Vargiu et al., 2016).

**Figure 2:**
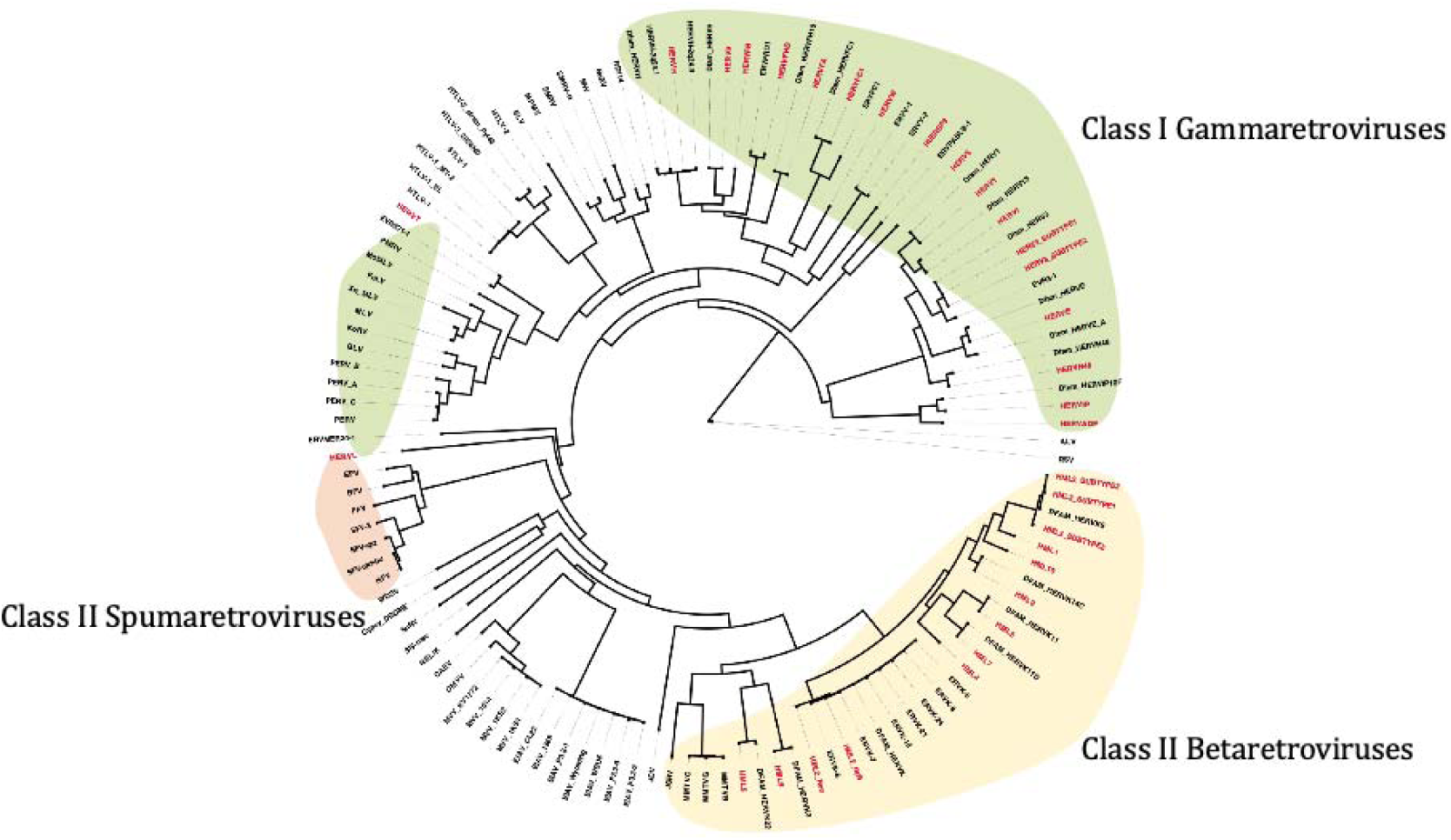
Phylogeny of the reconstructed Envs with the envelope protein of exogenous retroviruses. The phylogenetic tree was generated for the reconstructed HERV Env proteins with the 69 exogenous envelope proteins for Class I, Class II and Class III retroviruses. The class I exogenous and endogenous retroviruses are highlighted in green, class II in yellow and class III in orange color. The reconstructed Envs are presented in red color in the tree. The tree also contains RVs of other class with which no ERVs were clustered and hence are not highlighted.

### Distribution of ERVs in Primate Genomes

To track the spreading of the above HERV groups across primates’ evolution and evaluate their conservation, we used the above HERV Env consensus sequences to identify the *env* genes of class I, class II and class III ERVs through a group specific tBLASTn search in a the genome of 55 primate species. From the tBLASTn similarity search, we obtained 1374 hits for class I i.e. gamma-like *env*, 873 hits for class II i.e. beta-like *env* and 96 for class III i.e. spuma-like *env*. For all the three classes, we generated nt alignments and performed maximum likelihood (ML) phylogenies using IQTREE software with a bootstrap of 1000 replicates.

For the gamma-like *env* 1374 hits the best fit model generated for the phylogeny using the IQTREE was TVM+F+R4 (Figure 3a, Supplementary file 3). The phylogenetic analysis revealed group-specific clustering of env genes, primarily within the HERVHF supergroup, which includes HERVH, HERVFA, HERVFB, and HERVFC. The HERVHF supergroup is believed to have integrated into the *Catarrhini* lineage around 30–45 million years ago (Li et al., 2022). Our study also observed a widespread distribution of this supergroup, with most occurrences in Hominidae genomes, followed by other species. (Table 2, Figure 3a, Supplementary file 3). Similarly, we also observed a wide distribution of ERVT, ERVW, ERVE and ERVFRD followed by some small clusters of ERVH48, ERV9 and ERVIP. Additionally, we noticed distinct occurrences and clusters of HERV15, along with HERVR, ERVV1, ERVV2 and PABL, despite their exclusion from the blast analysis. Although HERV15, HERVR, ERVV1, ERVV2 and PABL were excluded from the primary Blast analysis, we observed distinct occurrence and clusters of these sequences in the subsequent phylogenetic analysis. This is likely because the Env consensus sequences, even though being group specific, share regions of homology with the excluded sequences. As a result, the consensus sequences effectively captured related members of the supergroup, even those not explicitly included in the initial dataset (Figure 3a). Of note, the consistent group specific clustering of the reconstructed Env sequences observed in the phylogenetic trees supports the robustness of the consensus sequences in representing the broader supergroup.

**Table 2:**
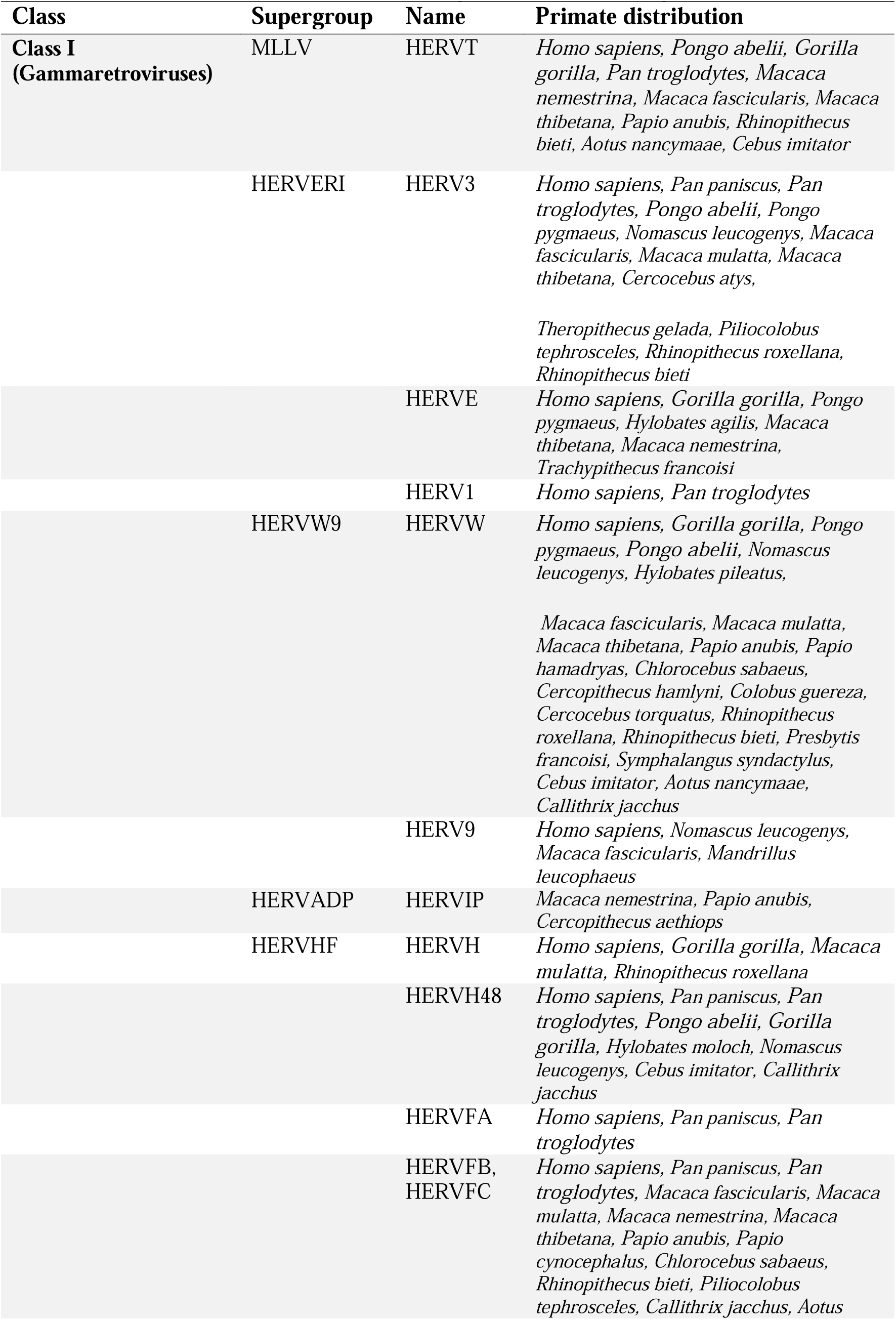
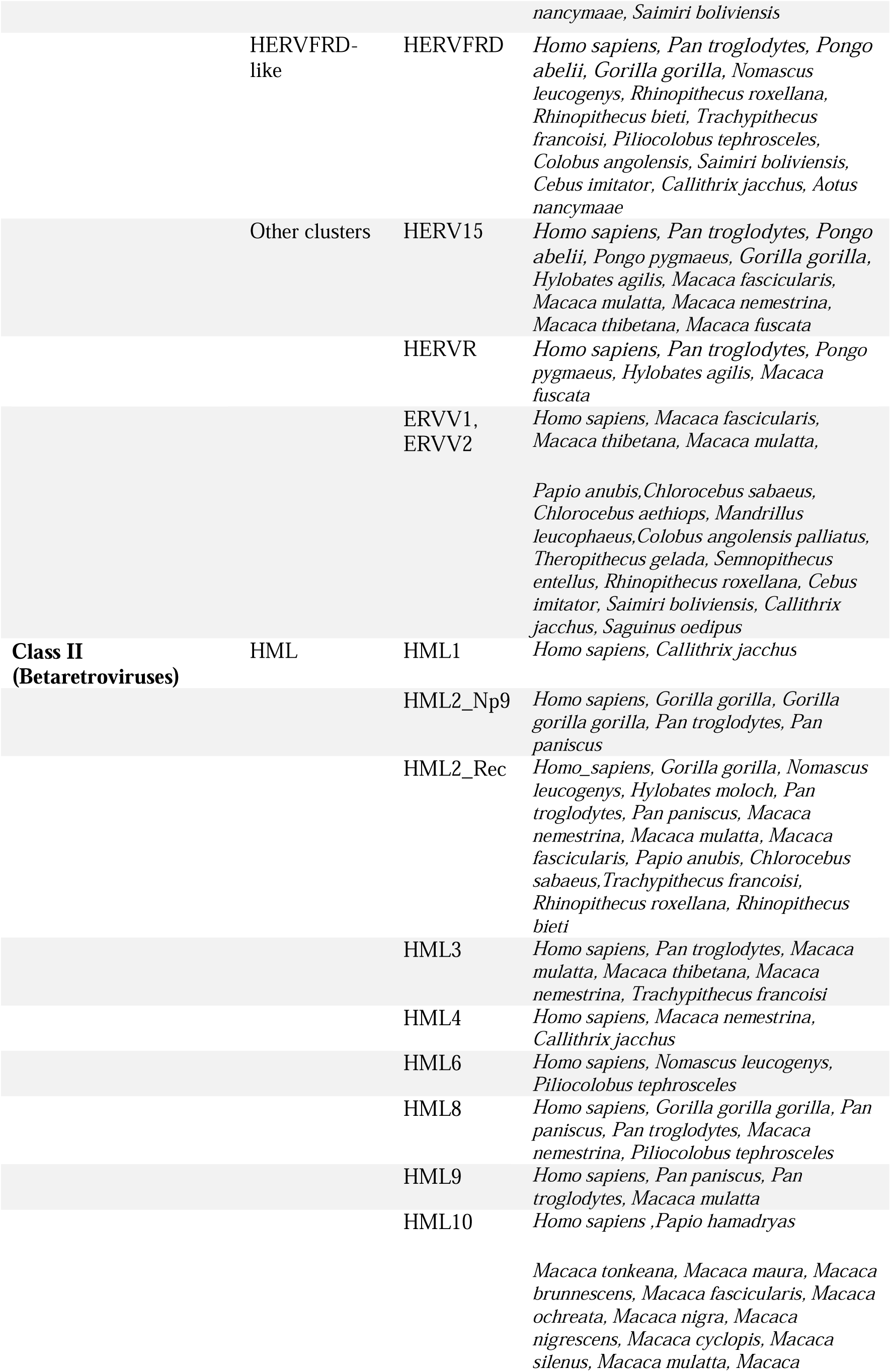
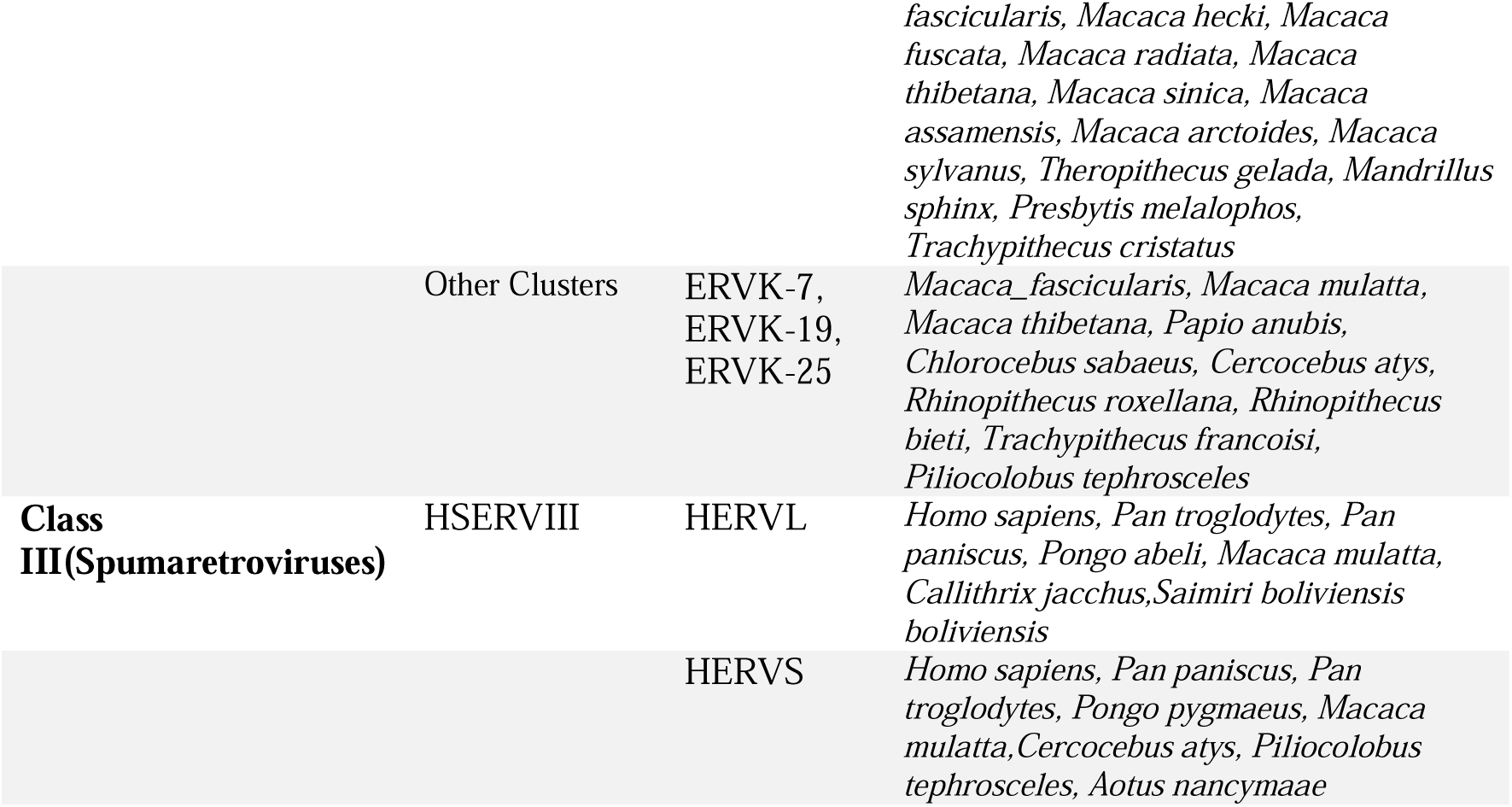
Widespread Distribution of ERV *env* genes across primate genomes.

Likewise, phylogenetic analysis was also performed for 873 beta-like *env* hits obtained from tBLASTn. The best fit model for the phylogeny obtained was TVM+F+I+R3, with the major grouping obtained for HML2, which was further divided into two subclusters corresponding to the presence of Np9 (type I) and Rec (type II) *env* splicing variants (Table 2, Figure 3b, Supplementary file 4). The HML2 group is present in abundance followed by HML8 and HML10 while smaller clusters were observed for HML1, HML3, HML6 and HML9 (Table 2, Figure 3b, Supplementary file 4). A very few sequences were obtained in the *Callithrix jacchus* that belong to *Platyrrhini* parvorder. In addition to the above clusters for HML2 Np9 and Rec, a third cluster of HML2 Rec included sequences that are annotated as ERVK-7, ERVK-19 and ERVK-25. The tBLASTn search for HML4 (HERVK-13) in the platyrrhini genomes, we obtained a fewer hits in *Callithrix jacchus,* and hence to confirm the presence of HML4 group in *Callithrix jacchus* and other Platyrrhini genomes we performed the BLAT search in two *Platyrrhini* genomes available on UCSC Genome Browser i.e. Marmoset (*Callithrix jacchus*; Callithrix_jacchus_cj1700_1.1/calJ) and Squirrel monkey (*Saimiri boliviensis*; Broad/saiBol1). We also performed the BLAT search for other HML supergroup members which have not been previously reported in the Platyrrhini parvorder.

From the BLAT search, we obtained 6 copies of HML2 annotated as HERVK in the squirrel monkey genome while no HML2 sequences were detected in *Callithrix jacchus* (Table 3). We also identified the presence of other members of HERVK supergroup i.e. HML1, HML4, HML8, HML9 and HML10 in the squirrel monkey genome that has not been reported previously (Table 3). The sequences identified are in few cases full-length ERV copies while in most of the cases are of very short length or fragmented. These results indicates two possible hypotheses for the presence of HML1, HML2, HML4, HML8, HML9 and HML10 in squirrel money: either the onset of these HMLs began even before 30 mya i.e. before the split between NWM and OWM, that occurred around 43 mya or they got fixed in to the genome as a results of cross-species transmission of RVs after the split.

**Table 3:**
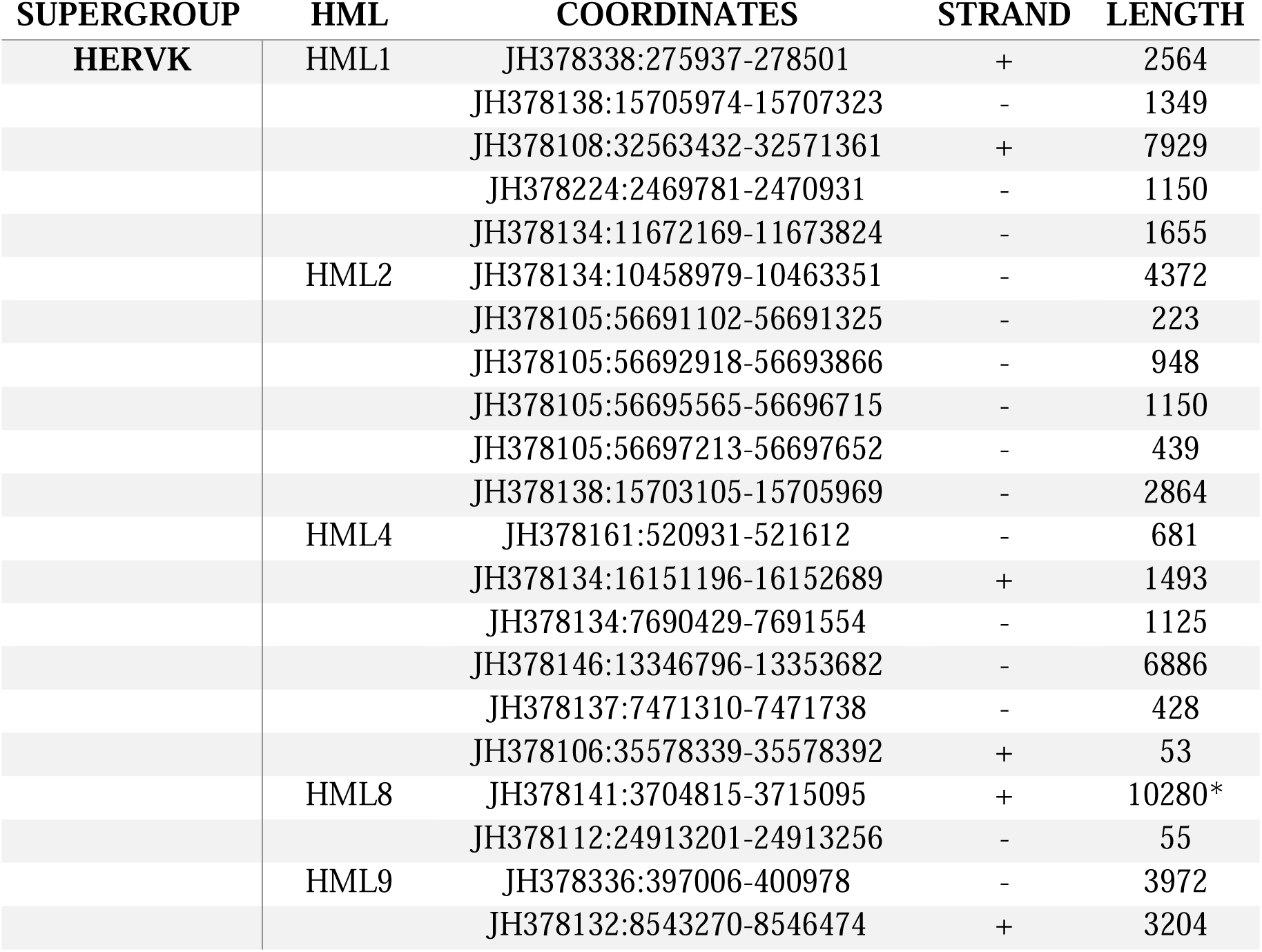
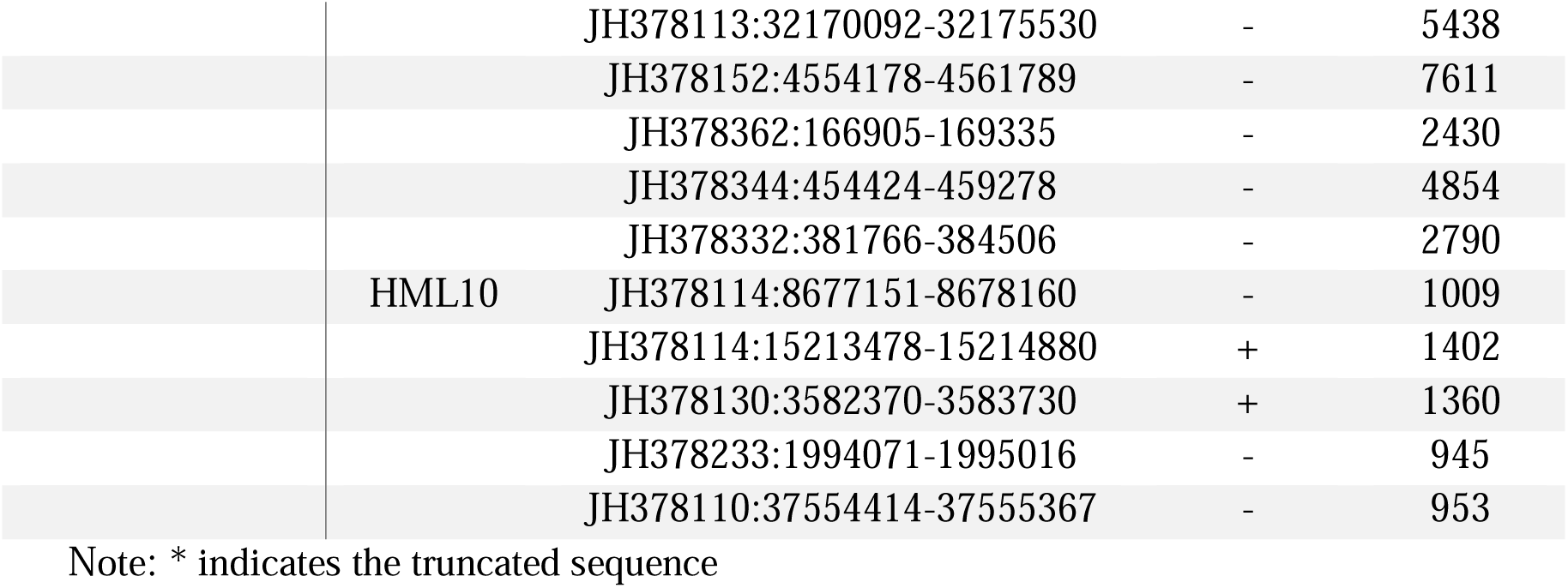
Coordinates of HML supergroup in *Platyrrhini* (Squirrel Monkey Genome)

While performing tBLASTn search for both HERVL and HERVS, hits were mostly obtained for gamma-like ERVs, specifically for ERV3-1, ERVFC-1, ERVFRD and ERV-W. These results were in line with our previous studies which states that HERVL is more similar to ERV3-1 in hyrax, tenrec, armadillo, alligator and turtle (Bénit et al., 1997; Cordonnier et al., 1995; Vargiu et al., 2016). The hits were further refined by excluding all the gamma-like ERVs as well as refined by excluding short elements and removing the sequence of other repetitive elements acquired by secondary integrations. Hence, we obtained only 96 hits specific to HERVL and HERVS. An ML phylogenetic tree with the best fit model of K3Pu+F+R3 was generated with the bootstrap of 1000 replicates (Figure 3c, Supplementary file 5). The phylogenetic analysis performed for these groups showed that a major distribution took place in *Homo sapiens* followed by *Macaca mulatta* and other primate genomes.

Apart from the group specific *env* clustering, we also observed some small clusters hinting towards the presence of recombinant ERVs, for example, the grouping of HML4 for *Callithrix jacchus* with HML2 in the beta-like *env* phylogeny.

### Evidence of Recombination in the ERVs’ Env

An important factor for the diversity of the *env* gene are recombination events, since recombination is a ubiquitous process that may give rise to new variants of existing viruses, possibly also accounting for changes of the original tropism. Hence, we aimed to detect the potential recombination events in our gamma, beta and spuma-like *env* alignments. To this purpose, we used RDP4 software, which implements multiple built-in methods to detect recombination events, which are confirmed when four or more methods yield a positive signal. The software identified multiple recombination events in the gamma and beta alignment files generated in this study (Figure 4a). Since the spuma-like *env* alignment includes only two groups i.e. HERVL and HERVS, and no recombination among them was detected, the spuma-like *env* were aligned with gamma-like *env,* to check the presence of recombination among the two classes. The RDP4 software provides an overview of each recombination event, including the major parent (the sequence contributing the majority of the recombinant), the minor parent (the sequence contributing smaller portions to the recombinant), and the breakpoint position (the nt position where the recombination occurs). Using the detected recombination events and their specific breakpoint positions, we performed BLAT searches in the UCSC Genome Browser for all gamma-like, beta-like, and spuma-like *envs*. While performing the BLAT search, we considered all the primate genomes available in Genome Browser (n=15), also with the purpose of tracing back the initiation of these recombination events along primates’ evolution (Figure 4b). From the BLAT analysis of gamma-like, beta-like, and spuma-like *env* sequences, we observed a distinct *env*-LTR recombination pattern. In all cases, the minor parents identified by the RDP software were LTRs within the *env* sequences.

**Figure 3:**
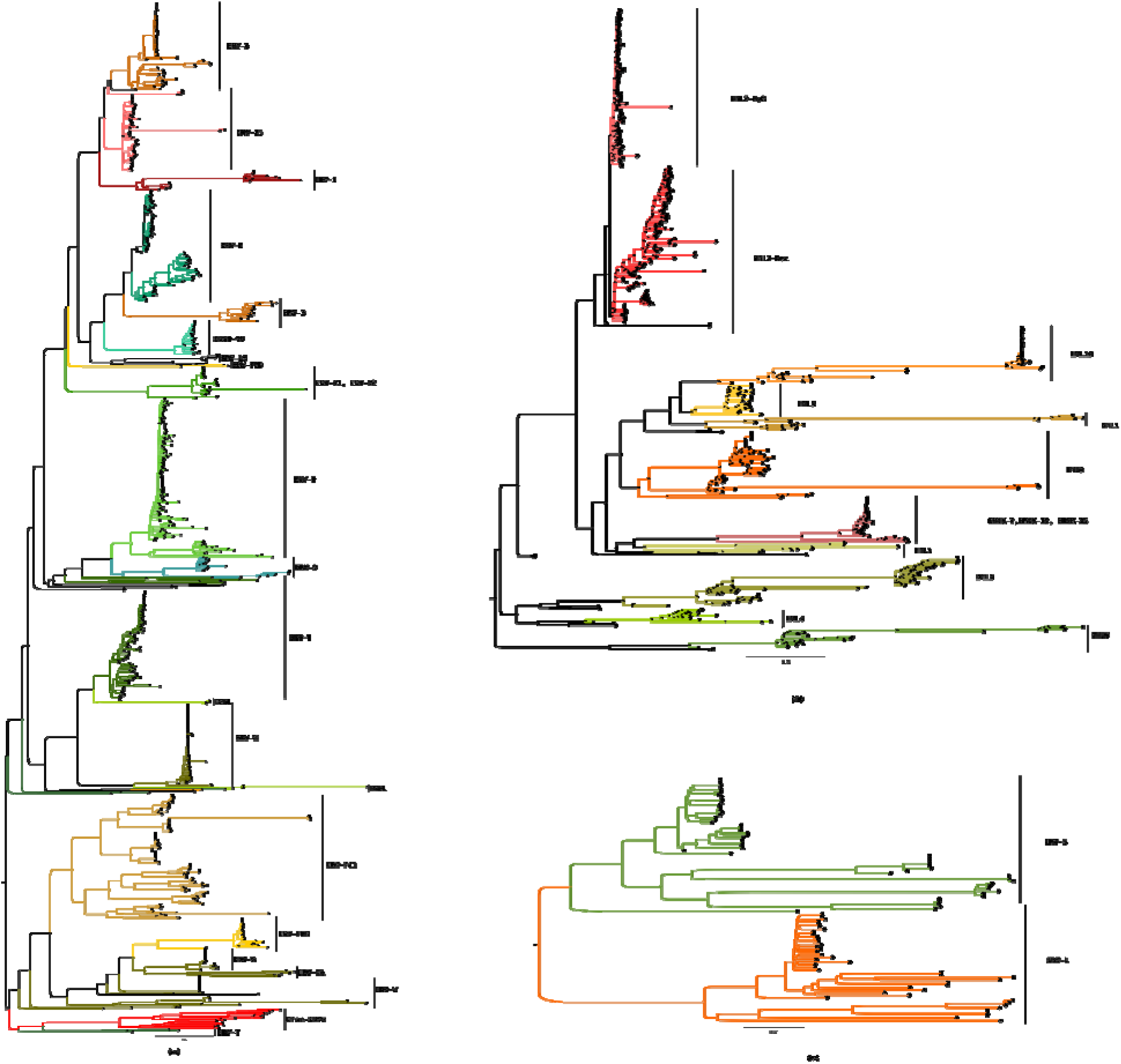
Phylogenetic analysis of the envelope hits obtained from the tBLASTn analysis for the three classes i.e. Class I (gamma-like *env*), Class II (beta-like *env*) and Class III (spuma-like *env*). The trees were generated using IQ-TREE software with the bootstrap of 1000 replicates. (a)Phylogeny for all the hits obtained for Class I ERVs with the best fit model of TVM+F+R4. (b) Phylogeny for all the hits obtained for Class II ERVs with the best fit model of TVM+F+I+R3 and (c) Phylogeny for all the hits obtained for Class III ERVs with the best fit model of K3Pu+F+R3. All the groups obtained from the tBLASTn hits and further clustered together with the phylogenetic analysis are labelled in the tree. Similar trees are provided as supplementary files with the tip labels having all the annotations and the accession numbers for all the three classes (Supplementary file 2,3 and 4).

**Figure 4:**
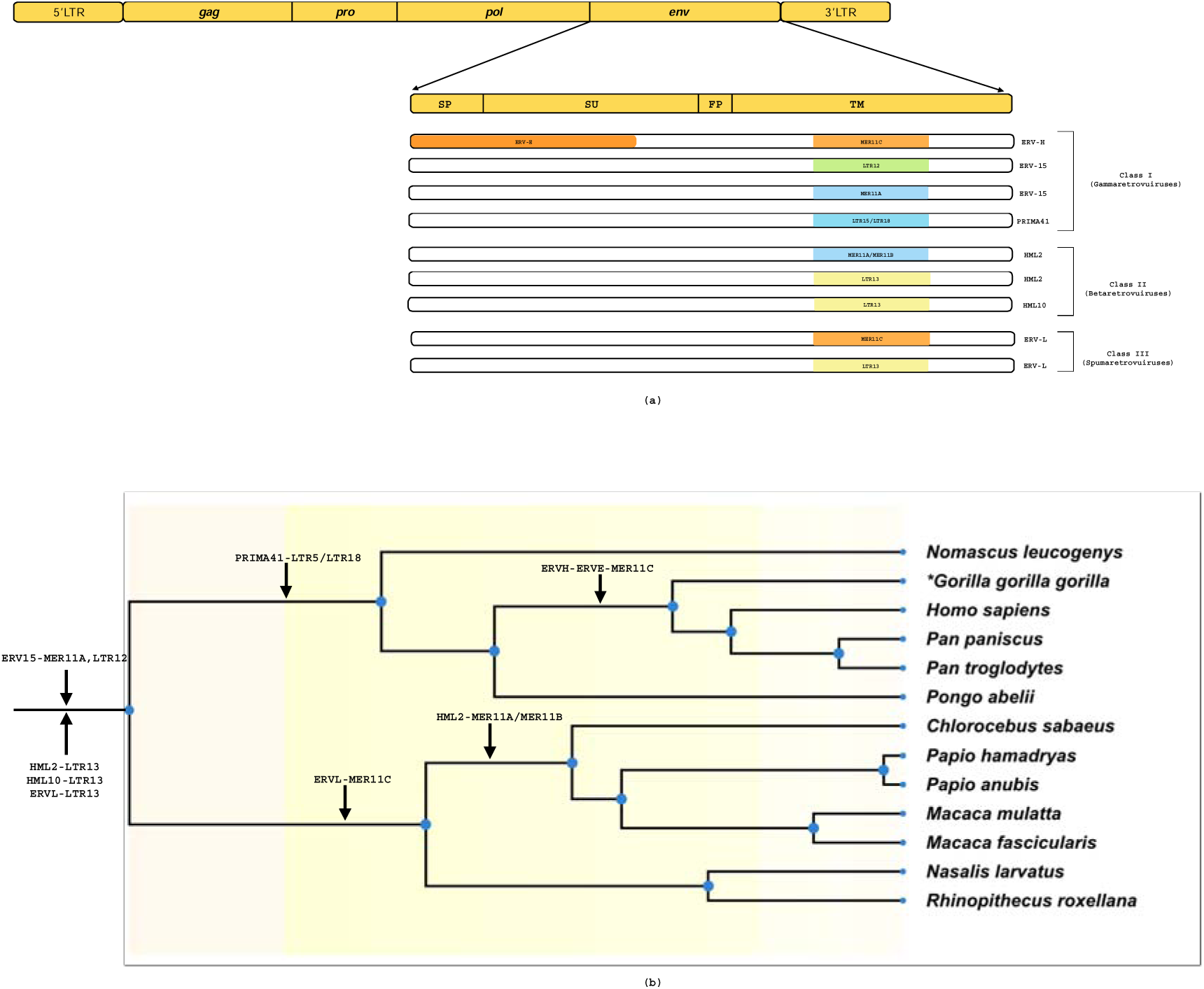
Detecting the recombination events in the ERV’s *env* gene. (a) Schematic representation of *env* recombination events in ERVs. The diagram indicates the location of the LTRs in the recombinant *env* gene, with the coordinates provided in supplementary file 5. (b) Phylogenetic tree illustrating the initiation of recombination events in the primate lineage. This tree was generated based on the available *Catarrhini* genomes in the Genome Browser using TimeTree server. Since the genome used by the server was *Gorilla gorilla gorilla* hence is marked with *. The presence of recombination events was tested using BLAT searches in the Genome Browser to trace the timing of their initiation and hence annotated on the phylogenetic tree based on the genomes the recombinants were detected.

The recombination events detected in the gamma-like *env* alignment are ERVH-ERVE- MER11C, ERV15-LTR12, ERV15-MER11A, and PRIMA41-LTR5/LTR18. The first event was detected in ERVH, in which ERVE was present in the SU region while MER11C that is an LTR of HML8 group was present in the TM region of the *env* (Figure 4a). This event was only identified in 4 genomes (Table 4, Supplementary file 6), suggesting a very recent occurrence (i.e. less than 9 mya) (Figure 4b). In fact, the recombinant sequences were shared only between 4 species of *Catarrhini* parvorder and were absent in more distantly related primates (such as gibbons, orangutans, etc.). This observation, together with the presence of shared recombination sites, suggest that the insertion occurred after the divergence of these species, while its absence in others indicate that it is a recent recombination event. It can be said that this recombinant form of ERVH emerged possibly due to the co-packaging of ERVH, ERVE and HML8 and thus the recombinant form ERVH-ERVE-MER11C was fixed in the genome of human, chimpanzee, gorilla, and bonobo with the very few copies. Other recombination events were observed in ERV15 and PRIMA41, in which a LTR was detected in the TM region of the *env* gene. The ERV15-LTR12/MER11A was detected in 12 primate genomes while the PRIMA41-LTR15/LTR18 that involves the LTR of ERV15 and ERV3 groups, respectively, was observed in only 6 genomes (Table 4, Figure 4b). Of note, these two events were present as a single copy in each genome in just one chromosome, as indicated in supplementary file 6. These single events might have occurred due to the template switching mechanism during the reverse transcription process and eventually were fixed in the chromosome.

Similar events were detected in beta-like *env* alignment, where the LTRs were detected in the TM region of the HML *env* gene. The events detected were HML10-LTR13, HML2-LTR13 and HML2-MER11A/MER11B (Figure 4a). Single chromosomal recombination event was detected for HML10-LTR13 and HML2-LTR13, where the presence of LTR13, which is the LTR of the HML4 group, was detected in the HML10 and HML2 *env* gene in 9 genomes (Table 4, Figure 4b). Similar to ERV15, these events are also present as single copies in the primate genomes. Unlike the above mentioned single chromosomal recombination events, the HML2-MER11A/MER11B recombination event was found in high abundance in the genomes of species from the *Cercopithecinae* and *Colobinae* subfamilies. However, within the *Hominidae* family, this recombination was detected in only a single species, *Nomascus leucogenys*. Especially, abundant number of copies of this recombinant were detected in the two Macaque genomes i.e. *Macaca fascicularis* and *Macaca mulatta*. The HML8 group contains three types of LTR i.e. MER11A, MER11B and MER11C (Scognamiglio et al., 2023), of which the presence of MER11A in HML2 *env* was detected in *Chlorosebus sabaeus, Macaca fascicularis, Macaca mulatta, Papio hamadryas, Nasalis larvatus* and *Rhinopithecus roxellana* while MER11B was detected in *Nomascus leucogenys, Chlorocebus sabaeus, and Macaca fascicularis* (Table 4, Figure 4b) suggesting that the HML2 template switching took place with two of the three above different variants of HML8. Since no evidence of this recombination event was found in most of the species of *Hominidae* except for *Nomascus leucogenys,* it is likely that it occurred after the split between *Colobinae* subfamily and *Platyrrhini*, simultaneously with the abundant spread of HML2. All the recombinations detected in the HML group are hypothesized to be emerged due to the template switching mechanism as in the gamma-like HERVs and hence were fixed in the primate genomes.

Lastly for the spuma-like *env,* we detected two recombination events for ERVL i.e. ERVL- MER11C and ERVL-LTR13 (Table 4, Figure 4a). These two single chromosome recombination events were detected in six species genomes (Table 4). As mentioned for all the above recombination events, ERVL seems to have also followed the same mechanism of template switching for both the LTRs.

Finally, to infer the functional effect of the presence of LTRs in the *env* region of ERVs, we performed the conserved domain (CD) search in NCBI. We first used the reconstructed Env to identify the domains present in the *env* and then performed the same search with the recombinant forms. In Table 5, all the CDs are listed, reporting their position in the *env* obtained from NCBI for the non-recombinant forms as well as their presence or absence in the recombinant forms. Of note, a consistent pattern of loss of CD is observed for the ERVs which have an LTR in the *env* gene. The domains are either completely absent or truncated due to the recombination with the LTR, leading to the loss of function in the corresponding protein. As shown in Table 3, the only domain identified for ERVH is TLV coat, but other domains have been identified for the recombinant form of ERVH-ERVE-MER11A suggesting an *env* acquisition event for the ERVH group by the co-packaging with ERVE. From the overall results, it seems that the LTRs of HML8 (MER11A, MER11B and MER11C) are the most frequently found in the *env* gene of different ERVs and are associated with recombination in primate genomes.

**Table 4:**
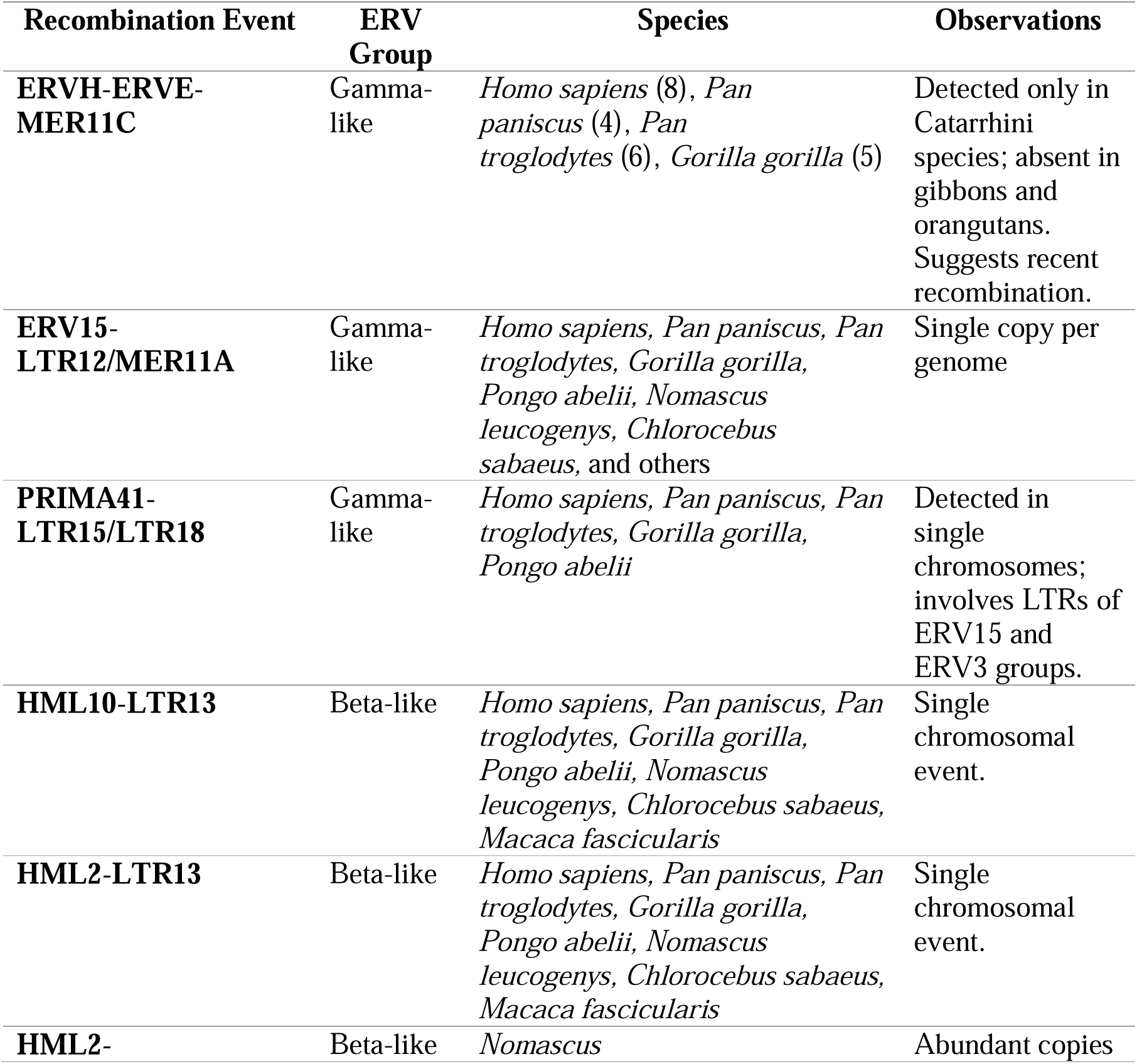
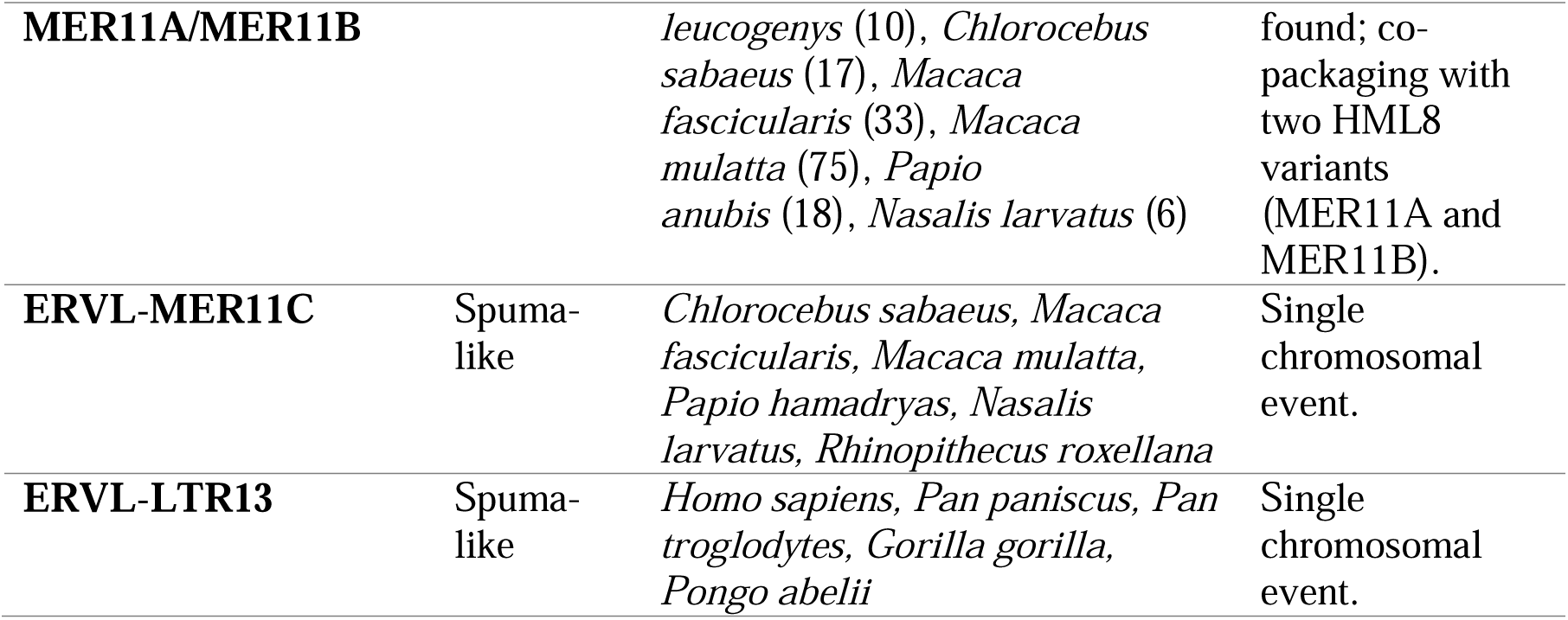
Summary of Recombination Events in Gamma-like, Beta-like, and Spuma-like Env Genes Across Primate Genomes.

**Table 5:** Detection of the conserved domains in the reference ERV sequences as well as recombinant sequences.

| Recombinant ERV | Conserved Domains Features |  |  |  |  |  |  |  |  |
| --- | --- | --- | --- | --- | --- | --- | --- | --- | --- |
|  | HERV-K_REC Super family | HERVK_en v_2 | TLV Coat | Ebola-like_HR1-HR2 Superfamily |  |  |  |  |  |
|  |  |  |  | HR1-GP1 interface | HR2 | Homotrimer interface | C1 binding site | CX(6,7)C Motif | Immuno-suppressive region |
| **\*ERV15** | - | - | 1064-1258 | 1439-1552 | 1631-1669 | 1439-1669 | 1517-1528 | 1571-1598 | 1521-1569 |
| **ERV15-LTR12** | - | - | - | √ | - | √* | - | - | √ |
| **ERV15-MER11A** | - | - | - | - | - | - | - | - | - |
| **\*ERVH** | - | - | 825-1487 | - | - | - | - | - | - |
| **ERVH-ERVE-MER11C** | - | - | - | √ | - | √ | √ | √ | √ |
| **\*PRIMA41** | - | - | - | - | - | - | - | - | 929-955 |
| **PRIMA41-LTR5/LTR18** | - | - | - | - | - | - | - | - | - |
| **\*HML2** | 55-315 | 346-849 | - | 1501-1659 | 1753-1878 | 1507-1878 | - | - | - |
| **HML2-LTR13** |  | - | - | - | - | - | - | - | - |
| **HML2-MER11A/MER11B** | √ | √ |  | √ | - | √* | - | - | - |
| **\*HML10** |  | 157-636 | - | 1339-1437 | - | 1339-1497 | - | - | 1421-1447 |
| **HML10-LTR13** |  | - | - | - | - | - | - | - | - |
| **\*ERVL** | - | - | 455-1051 | 716-814 | 908-934 | 716-934 | - | - | 797-823 |
| **ERVL-MER11C** | - | - | - | - | - | - | - | - | - |
| **ERVL-LTR13** | - | - | - | - | - | - | - | - | - |

Note: √ indicates presence of the Domain; √\* indicates the domain is truncated; - indicates absence of the domain; \* indicates the non- recombinant ERV

## Discussion

ERVs are “fossils” of the ancient exogenous RVs that have become extinct, that can still carry out certain functional roles (Sverdlov, 2000). Indeed, the genomic integration of ERVs and its widespread dispersion has led to the domestication of some ERV proteins for useful functions. The best known examples of the conservation of an open reading frame (ORF) linked to the expression in appropriate tissue are syncytin-1 and syncytin-2 encoded by HERVW (ERVWE1) (Blond et al., 2000; Grandi & Tramontano, 2017) and HERVFRD-1 (ERVFRD-1) (Rote et al., 2004), respectively. Another recurrent ERV domestication is aimed to its use as a restriction factor and interfering in steps of exogenous RV infection (Frank & Feschotte, 2017). Available ERV Env studies are mostly focused on finding their possible role in physiological or pathological conditions, but limited studies have been performed to understand ERVs’ evolutionary dynamics. For this, a focus on how various factors may affect the Env divergence is required.

The initial step in understanding such ERVs evolutionary dynamics – which encompasses their origin, proliferation within the host genome, and the mechanisms of inactivation and elimination – is to precisely reconstruct their viral genes that can also be essential to define their host-virus interactions. In addition, understanding their residual functions can help both in elucidating their potential role in various pathological conditions, and allow their exploitation as antivirals, gene therapy vectors, etc. In this scenario, an Env reconstruction clearly aids in broadening the understanding and discovering of various aspects of currently existing ERVs (Johnson, 2019). Hence, with the intention of studying the ERV Envs divergence, we firstly reconstructed 32 ancestral ERVs Env prototypes that can help in various functional analysis such as gene regulation and expression, pathogenicity, evolutionary dynamics, host-ERV interaction, etc. (Johnson, 2019). The screening of 55 primate genomes in *Catarrhini* (*Hominidae, Cercopithecinae, Colobinae*) and *Platyrrhini* parvorders using the reconstructed Env sequences for each HERV class demonstrated that the majority of ERV *env* nt sequences hits are obtained in the *Catarrhini* parvorder specifically in Hominidae Superfamily. In the phylogenetic analyses performed for each Class, we observed a group-specific clustering of all the ERVs. Over time, the genomic persistence of ERVs has been accompanied by insertions, deletions, mutations, and even the loss of the *env* gene, all of which contribute to biases in their phylogenetic analyses. As a result, we report some mixed clustering among the ERV classes. Apart from the 17 gamma-like *env*, we also observe hits for ERV15, HERVR, ERVV1, ERVV2 and PABL *env* genes which were initially not included in the dataset, meaning that the generated consensus sequences represent very well the members of the same supergroup. Our investigation demonstrates that the superfamilies HERVHF and HERVW9 of gamma-like *env*, HML2 group of beta-like *env* integrated in the primate lineage over 30-45 mya, earlier than previously reported (Li et al., 2022). In fact, several studies focused on the characterization and identification of HERVs in the primate genome - also reported the presence of the HML supergroup in the *Catarrhini* parvorder (Grandi et al., 2017, 2021; Scognamiglio et al., 2023) and by far HML5 has been known to be the most ancient member of the HERVK supergroup (Lavie et al., 2004), as its presence has been previously reported in marmoset and squirrel monkey. These reports indicate that their integration began after the split between *Catarrhini* and *Platyrrhini* i.e. ∼30 mya, as the HML sequences were not detected in *Callithrix jacchus* genome except for HML5. However, we identified the presence of members of HERVK supergroup i.e. HML1, HML2, HML4, HML8, HML9 and HML10 in the squirrel monkey genome that has never been previously reported, while no HML2 sequences were detected in *Callithrix jacchus* (Table 3). Thus, we show that the onset of integration of HML1, HML2, HML4, HML8, HML9 and HML10 initiated in the *Platyrrhini* even before 43 mya, possibly having a lower rate of amplification as they are less numerous in this parvorder, while they were spread more abundantly after the split of *Catarrhini* and *Platyrrhini*. Differently from gamma-like and beta-like ERVs, we detected only 96 hits for spuma-like *env.* the reason for such lower prevalence of spuma-like ERVs in the genome could be that they either less frequently integrate into the germ line or might have a shorter persistence in the genome as compared to gamma-like or beta-like ERVs (Hayward et al., 2015). Overall, the present phylogenomic approach suggests the widespread presence of the gamma ERVs, indicating that they may have been integrated in the host genomes through multiple sources, and demonstrates the onset of class II ERV even before the split between *Catarrhini* and *Platyrrhini* parvorders. A complex evolutionary trajectory as compared to the one of spuma-like ERVs.

Numerous studies have reported multiple recombination events in ERVs in the vertebrate lineage leading to several modifications throughout the endogenization process (Chabukswar et al., 2023). Recombination mechanisms contributing to the observed variation in the ERVs primarily involve template switching during reverse transcription. Other processes, such as homologous recombination within the host genome, can also lead to the exchange of genetic material between ERVs. While cross-species transmission and *env* acquisition or degradation are critical processes in shaping ERV evolution and diversity, they are not direct mechanism of recombination but rather represent the outcome or influences on recombination (Chabukswar et al., 2023; Pérez-Losada et al., 2015). Such events can greatly increase the diversity among the ERVs and complicate their classification. Hence, to detect the presence of such recombination events in primates we performed recombination analysis and identified 8 potential recombination events i.e. ERVH-ERVE-MER11C, ERV-15-LTR12, ERV-15- MER11A, HML2-LTR13, HML2-MER11A, HML2-MER11B, HML10-LTR13, ERVL-MER11C and ERVL-LTR13 in gamma-like and beta-like and spuma-like *env* sequences (Table 4). We confirmed these events using the genome assemblies available in Genome Browser, hence assuming the presence of these recombinants in the other primate genomes of the superfamily.

The potential mechanism due to which such events occur in the genomes is the co-packaging of two different ERVs or an ERV and an exogenous RV in a virus particle, followed by the template switch between the two genomes. In the template switching mechanism the two RV copies are co-packaged into the virus particle, and the template switch occurs as the RT dissociates from one template to another resulting in intermolecular i.e. homologous or nonhomologous recombination or intramolecular i.e. deletions or insertions (Delviks-Frankenberry et al., 2011). As in the case of the recombinants detected in the present study, a template switch could have occurred between the highly conserved TM region of the *env* of one ERV and the LTR of the other ERV due to sequence similarities, resulting in the observed recombination of an LTR inserted into the TM region of the *env* gene. According to the mechanism of template switching, the LTR found within the *env* gene has the features of a viral RNA genome LTR (i.e. not with the complete U5-R-U3 structure that characterizes the proviral ones), as indeed it is observed in the sequences. Additionally, it is possible that the observed recombinant events arose from the co-packaging from ERV and a related exogenous RV, especially if the latters were still actively circulating at the time when the events occurred. Such interactions could have facilitated inter-viral recombination, leading to the integration of recombinant sequences into the host genome. A prominent example of such event is the recombination between exogenous MuLv and its endogenous form leading to the generation of recombinant polytrophic MuLVs that utilize a different cell surface receptor for their entry, altering the host range and increasing their pathogenic potential (Alamgir et al., 2005; Chabukswar et al., 2023; Evans et al., 2009, Williams et al., 2024).

As in case of ERVH-ERVE-MER11C, ERVH seems to acquire the *env* gene of the ERVE group, leading to the generation of a recombinant ERV that retains the potentially coding ERVE Env, while for the rest of the recombinant sequences an LTR exchange with the TM region of the *env gene* is observed, eventually leading to loss of the *env* function, further promoting the intragenomic spreading of *env*-less ERVs (Magiorkinis et al., 2012). Interestingly, such swapping of the *env* region can also possibly lead to the emergence of different variants of the existing RV, as in case of HML2-MER11A/MER11B, where it is predicted that different HML8 variants swapped their LTR with the TM region of the HML2 *env*, and thus spreading in abundance in the *Cercopithecinae* subfamily members and creating diversity in the HML2 group. The recombination between HML2-MER11A/B in the macaque genomes was previously reported (Chabukswar et al., 2022; Williams et al., 2024), suggesting that the HML8 region appeared in HML2 *env* region due to the recombination between HML2 and HML8 virus due to co-packaging of HML2 and HML8 RNAs during the reverse transcription and has been experimentally verified by Williams et al., 2024. Similar to HML2-MER11A/B recombination, various recombinant events have been reported in this study for gamma-, beta- and spuma-like Envs, having the LTRs in the TM region of the *env.* Hence, as suggested, it is worth considering that these events can have implications in the biology of these viruses (Williams et al., 2024). These kind of events have also been previously reported for Harlequin which have a complicated structure of LTR2-HERVE- MER57I-LTR8-MER4I-HERVI-HERVE-LTR2 (Vargiu et al., 2016). The recombination events described in the present study provide more insights of the prevalence of recombinant ERV forms within the host genome, underlying the role of recombination as a driving force behind the ERVs diversification.

## Conclusion

ERVs are the remnants of ancestral exogenous RVs whose origin and diversity are still not completely understood and are a great tool to study the coevolution of host and RVs. The present study provides a phylogenomic perspective of the retroviral *env* gene diversity. With this study approach we provide the evidence for inter- as well as intra-group recombination events. Present results demonstrate the prevalence of ERVs across the primate lineage as well as the diversification of the retroviral Env through the process of recombination over the evolutionary timescale. Overall, this analysis gives further knowledge in studying the host-virus interactions on a larger scale.

## Supplementary Files

**Supplementary File1:** Excel sheet for all the HERVs used for the Reconstruction of HERVs’ Env (S1, S2 and S3). The S4 sheet contains Fasta File of gamma-like, beta-like and spuma-like reconstructed HERV Envelope sequences.

**Supplementary File2:** List of the primates’ genomes and the genome aseemblies that were screened during the tBLASTn search for screening the ERVs’ Env

**Supplementary File3:** Complete phylogenetic tree with the tip labels for Gamma-like ERVs’ env.

**Supplementary File4:** Complete phylogenetic tree with the tip labels for Beta-like ERVs’ env.

**Supplementary File5:** Complete phylogenetic tree with the tip labels for Spuma-like ERVs’ env.

**Supplementary File6:** Excel file for the Coordinates of the recombinant sequences in the 15 genome assemblies extracted from the BLAT search.

## Data availability statement

The original contributions presented in the study are included in the article/Supplementary material, further inquiries can be directed to the corresponding author.

## Author contributions

SC performed the analysis and wrote the manuscript. ES participated in performing the analysis. NG supervised the analyses, checked the data, and participated in editing the manuscript. ET and LTL conceived and coordinated the study. All authors contributed to the article and approved the submitted version.

