## Supplementary File 2 for "Screening Envelope Genes Across Primate Genomes Reveals Evolution and Diversity Patterns of Endogenous Retroviruses"

**Supplementary File 2:** List of the primates’ genomes and the genome aseemblies that were screened during the tBLASTn search for screening the ERVs’ Env

| No. | Primate | Genome Assembly |
| --- | --- | --- |
| 1 | *Homo sapiens* | GCF_000001405.40; GCF_009914755.1 |
| 2 | *Gorilla gorilla* | - |
| 3 | *Gorilla gorilla gorilla* | GCF_029281585.2 |
| 4 | *Pan troglodytes* | GCF_028858775.2 |
| 5 | *Pan paniscus* | GCF_029289425.2 |
| 6 | *Pongo abelii* | GCF_028885655.2 |
| 7 | *Pongo pygmaeus* | GCF_028885625.2 |
| 8 | *Nomascus leucogenys* | GCF_006542625.1 |
| 9 | *Hylobates agilis* | GCA_963574095.1 |
| 10 | *Hylobates pileatus* | GCA_021498465.1 |
| 11 | *Hylobates moloch* | GCF_009828535.3 |
| 12 | *Macaca fascicularis* | GCF_037993035.1 |
| 13 | *Macaca mulatta* | GCF_003339765.1 |
| 14 | *Macaca nemestrina* | GCF_000956065.1 |
| 15 | *Macaca thibetana* | GCF_024542745.1 |
| 16 | *Macaca fuscata* | GCA_003118495.1 |
| 17 | *Macaca tonkeana* | GCA_963573625.1 |
| 18 | *Macaca maura* | GCA_963574855.1 |
| 19 | *Macaca brunnescens* | - |
| 20 | *Macaca ochreata* | - |
| 21 | *Macaca nigra* | GCA_928851695.1 |
| 22 | *Macaca nigrescens* | - |
| 23 | *Macaca cyclopis* | GCA_026956025.1 |
| 24 | *Macaca silenus* | GCA_023807365.1 |
| 25 | *Macaca hecki* | - |
| 26 | *Macaca radiata* | GCA_963573735.1 |
| 27 | *Macaca sinica* | - |
| 28 | *Macaca assamensis* | GCA_023783095.1 |
| 29 | *Macaca arctoides* | GCA_021188215.1 |
| 30 | *Macaca sylvanus* | GCA_023807365.1 |
| 31 | *Papio anubis* | GCF_008728515.1 |
| 32 | *Papio cynocephalus* | GCA_963574715.1 |
| 33 | *Papio hamadryas* | GCA_023781915.1 |
| 34 | *Chlorocebus sabaeus* | GCF_015252025.1 |
| 35 | *Chlorocebus aethiops* | GCA_023783515.1 |
| 36 | *Cercopithecus hamlyni* | GCA_963574815.1 |
| 37 | *Cercocebus atys* | GCF_000955945.1 |
| 38 | *Cercocebus torquatus* | GCA_963574135.1 |
| 39 | *Colobus angolensis* | GCF_000951035.1 |
| 40 | *Colobus guereza* | GCA_030247045.1 |
| 41 | *Mandrillus leucophaeus* | GCF_000951045.1 |
| 42 | *Mandrillus sphinx* | GCA_023783085.1 |
| 43 | *Theropithecus gelada* | GCF_003255815.1 |
| 44 | *Rhinopithecus roxellana* | GCF_007565055.1 |
| 45 | *Rhinopithecus bieti* | GCF_001698545.2 |
| 46 | *Presbytis melalophos* | GCA_963575215.1 |
| 47 | *Trachypithecus cristatus* | GCA_963574045.1 |
| 48 | *Trachypithecus francoisi* | GCF_009764315.1 |
| 49 | *Piliocolobus tephrosceles* | GCF_002776525.5 |
| 50 | *Symphalangus syndactylus* | GCF_028878055.3 |
| 51 | *Cebus imitator* | GCF_001604975.1 |
| 52 | *Callithrix jacchus* | GCF_011100555.1 |
| 53 | *Saimiri boliviensis boliviensis* | GCF_016699345.2 |
| 54 | *Saguinus oedipus* | GCA_031835075.1 |
| 55 | *Aotus nancymaae* | GCF_030222135.1 |
