## Supplementary figures and images for "Screening Envelope Genes Across Primate Genomes Reveals Evolution and Diversity Patterns of Endogenous Retroviruses"

### Supplementary File 3

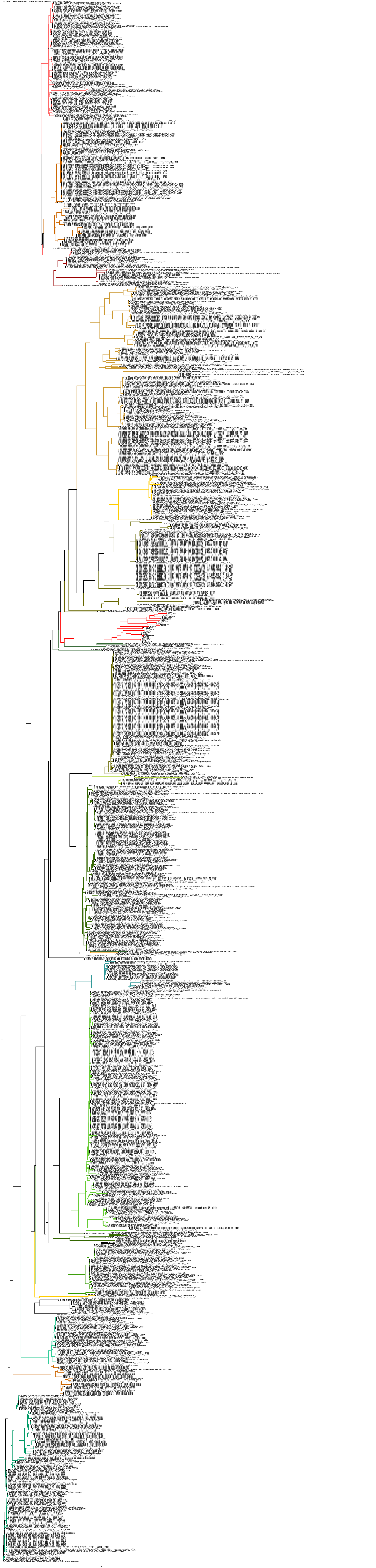

### Supplementary File 4

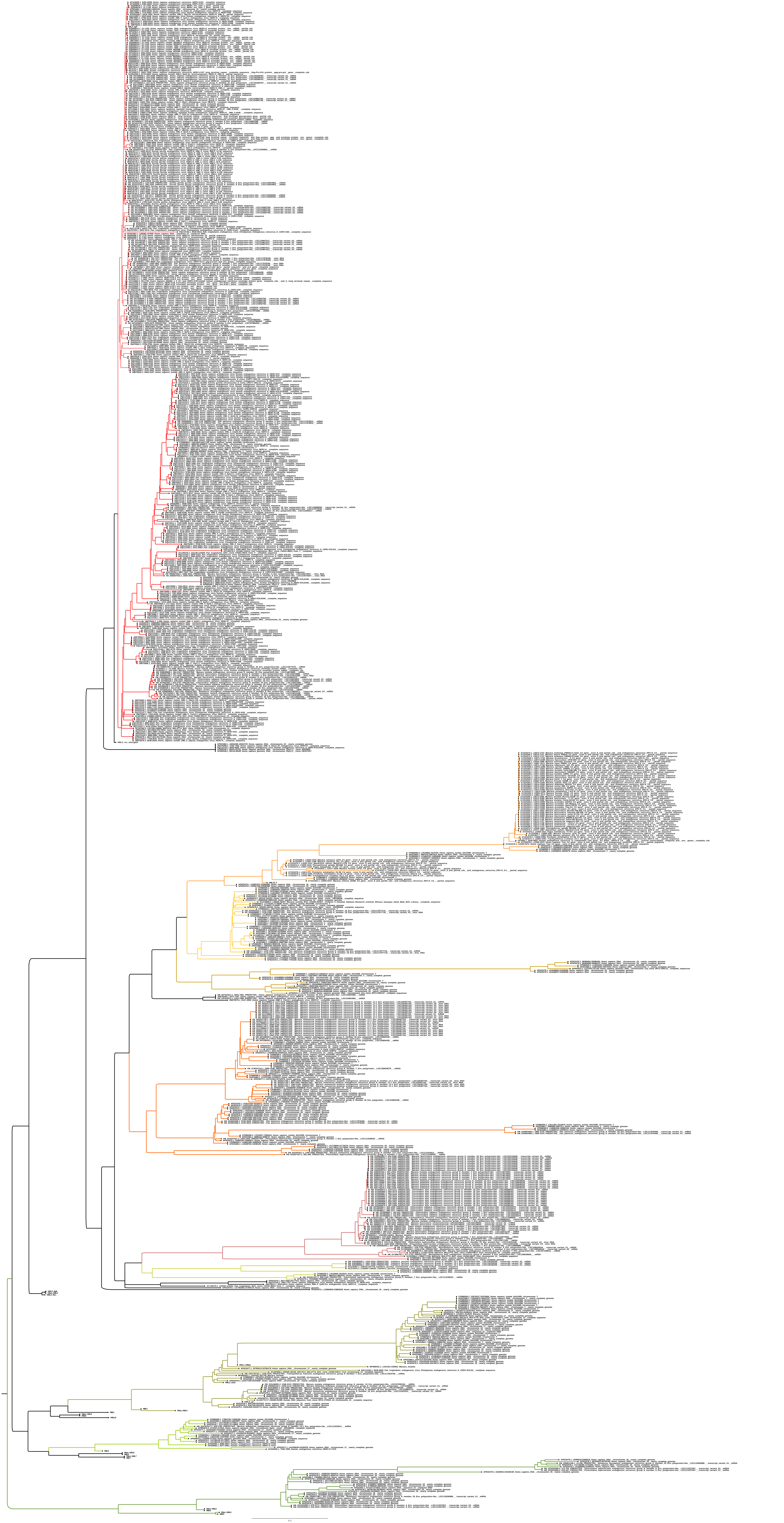

### Supplementary File 5

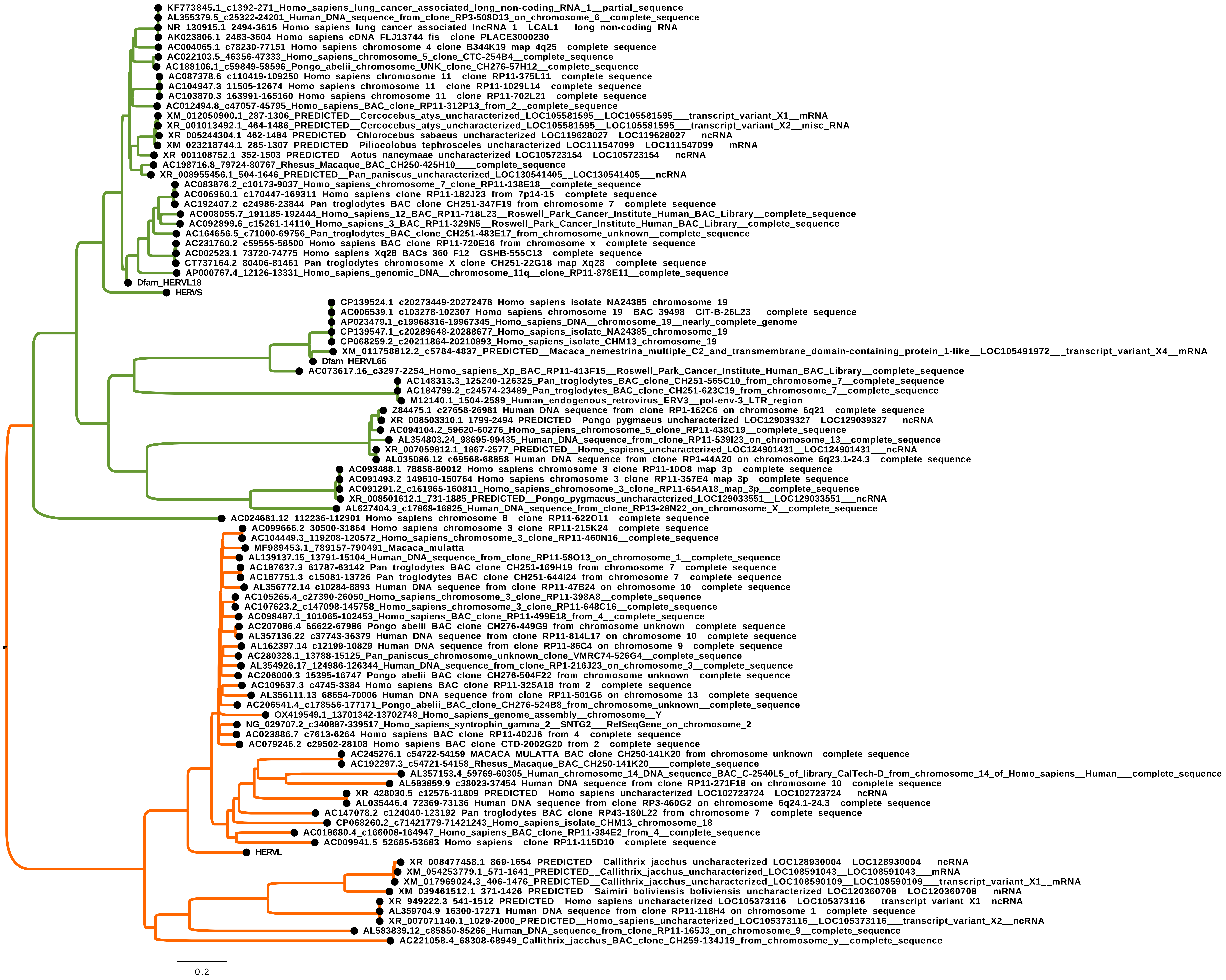
